# Short-lived alpha power suppression induced by low-intensity arrhythmic rTMS

**DOI:** 10.1101/2020.10.28.358986

**Authors:** Elina Zmeykina, Matthias Mittner, Walter Paulus, Zsolt Turi

## Abstract

This study was conducted to provide a better understanding of the role of electric field strength in the production of aftereffects in resting state scalp electroencephalography by repetitive transcranial magnetic stimulation (rTMS) in humans. We conducted two separate experiments in which we applied rTMS over the left parietal-occipital region. Prospective electric field simulation guided the choice of the individual stimulation intensities. In the main experiment, 16 participants received rhythmic and arrhythmic rTMS bursts at between ca. 20 and 50 mv/mm peak absolute electric field intensities. In the control experiment, another group of 16 participants received sham rTMS. To characterize the aftereffects, we estimated the alpha power (8-14 Hz) changes recorded in the inter-burst intervals, i.e., from 0.2 to 10 seconds after rTMS. We found aftereffects lasting up to two seconds after stimulation with ca. 35 mV/mm. Relative to baseline, alpha power was significantly reduced by the arrhythmic protocol, while there was no significant change with the rhythmic protocol. However, we found no significant long-term, i.e., up to 10-second, differences between the rhythmic and arrhythmic stimulation, or between the rhythmic and sham protocols. Weak arrhythmic rTMS induced short-lived alpha suppression during the inter-burst intervals.

## 1. Introduction

The self-organized activity of neurons and neural assemblies produces oscillating electric fields in the brain [1]. These oscillating electric fields are recurrent, as they feed back onto the neural assemblies thereby facilitating neural synchrony and plasticity [1]. Repetitive transcranial magnetic stimulation (rTMS) induces a periodic electromagnetic field in the brain [2], which triggers molecular, cellular, and electrophysiological changes in neuro-glia networks [3].

In our previous work, we studied the immediate electrophysiological effects of rTMS using a novel stimulation intensity selection approach [4]. In order to individually adapt the stimulation intensities, we prospectively estimated the rTMS-induced electric field strengths [4]. Using this approach we have shown that peak absolute electric fields between ca. 35 and 50 mV/mm already induced immediate changes in the electroencephalogram (EEG) in humans [4].

Yet, many applications of rTMS aim at inducing neural effects that outlast the duration of the stimulation itself. Therefore, in the present study we investigated possible aftereffects of the stimulation by focusing on the EEG recordings in the inter-burst intervals from 0.2 to 10 s after the rTMS bursts. The selected time window is free from rTMS-induced artifacts such as ringing, decay, cranial muscular, somatosensory or auditory artifacts [5].

To quantify the aftereffects, we estimated the spectral power in the alpha frequency band which is a common outcome measure in the rTMS-EEG literature [6]. Based on the *entrainment echo* hypothesis [7], we expected that rhythmic rTMS at the individual alpha frequencies would entrain neural oscillations and increase alpha power due to facilitated spike-timing dependent plasticity. On the other hand, we expected that arrhythmic (active control) or sham (90° tilt) protocols would not entrain ongoing posterior alpha oscillation and, therefore, would not produce any aftereffects.

## 2. Methods

### 2.1. Secondary analysis

To test our hypotheses we performed a secondary analysis of our openly available rTMS-EEG dataset (https://github.com/ZsoltTuri/2019_rTMS-EEG). We reported the immediate electrophysiological effects elsewhere [4]. This dataset contains EEG recordings from two separate experiments (see point 2.5 for more details).

### 2.2. Participants

We included only neurologically healthy participants in the study [4]. For more details, see Table 1.

**Table. 1.** Participant information. ^a^We assessed the handedness laterality index with the Edinburgh Handedness Inventory [8].

|  | Main experiment | Control experiment |
| --- | --- | --- |
| Sample size (n) | 16 | 16 |
| Mean age ± SD (years) | 25.5 ± 3.2 | 23.9 ± 3.9 |
| Age range (years) | 21 to 32 | 20 to 34 |
| Number of women/men | 8/8 | 8/8 |
| Exclusion criteria assessed by | Self-reports and/or neurological examinations |  |
| Contraindications | None | None |
| Mean laterality index<sup>a</sup> ± SD | 78.4 ± 50.1 | 78.8 ± 31.6 |
| Laterality index range | -30 to 100 | 0 to 100 |

### 2.3. Ethics

The Ethics Committee of the University Medical Center Göttingen approved the investigation, the experimental protocols, and all methods used in the main and control experiment (application number: 35/7/17). We performed all the experiments under the relevant guidelines and regulations. All participants gave written informed consent before participation [4].

### 2.4. Head modeling and electric field estimation

We used a freely available open software package called Simulation of Non-invasive Brain Stimulation (SimNIBS, version 2.0.1) [9]. We used anatomical T1- and T2-weighted and diffusion-based magnetic resonance imaging data (MRI) to generate individualized, multi-compartment head models. The head models included the following compartments (corresponding conductivity values in [S/m]): scalp (0.465), bone (0.01), cerebrospinal fluid (1.654), gray matter (0.275) and white matter (0.126). For the gray and white matter compartments, we used anisotropic conductivity values using the volume-normalized method [10].

### 2.5. Experimental procedure and stimulation parameters

In the main experiment (n = 16), we performed prospective electric field modeling to individually adapt the stimulation intensities (see Fig 1A). Participants took part in three rTMS-EEG sessions separated by at least 48 hours. In each session, we applied rTMS at 20, 35, or 50 mV/mm peak absolute electric fields. These field values correspond to 9.5 ± 1.1%, 16.8 ± 2%, and 23.9 ± 2.5 % of the group-averaged device output. We refer to these sessions as Low, Medium, and High intensity conditions, respectively. For further details about the rTMS protocols, see Fig 1B (top).

**Fig 1.**
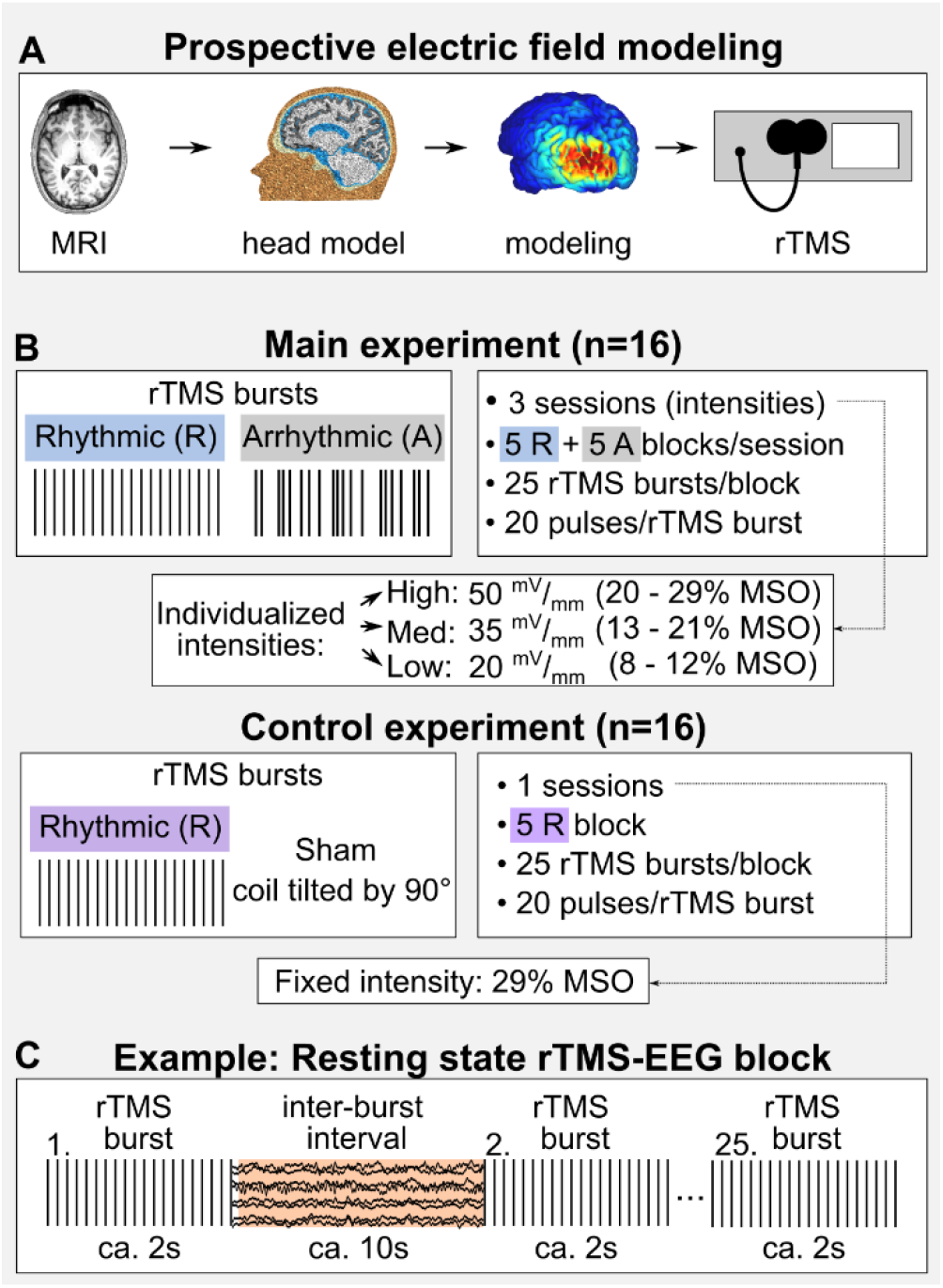
Study overview. (A) The stimulation intensity was individually adapted based on prospective electric field modeling. (B) The stimulation parameters in the main and control experiments. In the control experiment, we delivered rhythmic sham rTMS. (C) We defined the aftereffects by focusing on the rTMS artifact-free inter-burst intervals (highlighted in orange). Abbreviations: MSO – maximum stimulator output.

In the control experiment (n = 16), an independent group of participants received sham rTMS with the coil tilted by 90° (see Fig 1B, bottom) [11]. During the measurement, this sham protocol produced acoustic and ringing/decay artifacts while it minimized the induced electric field in the brain. We used the same stimulation intensity for each participant, which we fixed at 29% of the device output. This value corresponded to the maximum pulse amplitude used in the High intensity condition of the main experiment.

In both experiments, we applied rTMS over the left parietal-occipital area, specifically at the PO3 electrode as defined by the international 10/20 EEG system. The participants received the stimulation in the resting state, eyes open condition (Fig 1C). We delivered the rhythmic rTMS at the individual alpha frequency, which we estimated prior to each session from the resting state EEG recordings [4]. Based on the Arnold’s tongue model of neural entrainment, this is a necessary step to maximize the efficacy of inducing neural entrainment. In the arrhythmic rTMS, we applied rTMS in a manner that avoided any rhythmicity in the timing of the consecutive pulses [12,13]. Here, we prospectively adjusted the timing of each pulse so that frequencies in the alpha frequency band (8–12 Hz) as well as their harmonics and subharmonics did not occur (e.g., 4 and 16 Hz for 8 Hz) [4].

In both experiments, we used a MagPro X100 stimulator with MagOption (MagVenture, Denmark), normal coil current direction, biphasic pulses with 280 μs pulse width, and a MC-B70 figure-of-eight coil. During rTMS we simultaneously recorded the scalp EEG with a TMS-compatible, 64 channel, active EEG system (BrainProducts, Munich, Germany).

### 2.6. EEG analysis

#### EEG preprocessing

EEG analysis was performed using the FieldTrip software package (http://fieldtrip.fcdonders.nl) with custom-made MATLAB code. First, the TMS-EEG data were segmented into trials that were time-locked to the offset of the rTMS burst (from 3.5s before and 10 s after the last TMS pulse). The datasets in both experiments (main and control) included 125 trials with each stimulation condition. We removed the rTMS-induced ringing artifacts from 4 ms before to 9 ms after the TMS pulse. The first round of ICA (fastICA) was performed to automatically identify the decay artifact by averaging the time course of components over 50ms after each TMS pulse. Components with an amplitude exceeding 30 μV were rejected. Piecewise Cubic Hermite Interpolation (pchip) replaced the time intervals around the pulses.

Then, the data were downsampled to 625 Hz. We applied a 80 Hz low-pass and a 0.1 Hz high-pass filter (Butterworth IIR filter type, ‘but’ in FieldTrip). A discrete Fourier transform-based filter was used to remove the 50 Hz line noise. Next, the data were inspected for artifactual trials and channels. The procedure included a semi-automatic algorithm described in detail in reference [14]. In brief, we defined the outlier channels and trials, which exceeded 1.5 interquartile ranges. If a trial contained fewer than 20% of such channels, they were interpolated in the trial, but otherwise removed. The channels with line noise or high impedance levels were defined by estimating the correlation coefficient with the neighboring channels. We rejected channels that had a correlation coefficient value lower than 0.4 with their neighbors. All removed channels were then interpolated using the weighted signal of the neighboring channels.

After inspecting the data we defined the number of independent components for the ICA (binICA) by estimating the eigenvalues of the covariance matrix of the EEG data. We defined the number of ICA components as the rank of the diagonal matrix minus the number of the interpolated channels. We ran ICA only on trials that did not contain any interpolated channels. Independent components were visually inspected for artifacts.

The components containing eye-related artifacts, muscle, and line noise artifacts were projected out from the data. After preprocessing, 93.8±9.9 (mean ±SD) trials remained for the High, 91.1±13.4 trials for the Medium and 92.5±9.9 trials for the Low-intensity conditions. As the last preprocessing step, we applied two seconds of padding (‘mirror’) to the data intervals corresponding to baseline.

#### Short-term aftereffect

We performed the time-frequency analysis by running Wavelet decomposition on frequencies from 1 to 25 Hz for the whole length of the trial from −5.5 to 10 seconds around the TMS burst offset. The wavelet consisted of seven cycles with 3 Gaussian widths. Once the wavelet analysis was completed, we performed a statistical analysis to test the short-term aftereffect of the protocols and the time. To this aim, we used two-second intervals before (‘baseline’) and after (‘activation’) the rTMS burst. For each participant we averaged the data over all trials and then performed the statistical analysis (Fieldtrip as ‘actvsbslT’ test) separately for each intensity condition (High, Medium, and Low). To reduce the influence of the remaining TMS artifacts we performed a cluster-based permutation test (Monte Carlo, 2-25 Hz frequency range two-tailed t-test with 1,000 permutations) 0.2s after the last TMS pulse. The null hypothesis was rejected if the p-value of the maximum cluster level statistics was below 0.05 (one-tailed test).

#### Long-term after effect

For the second analysis, we normalized the power of all intervals of ca. 10 seconds length after rTMS bursts to baseline, i.e., the 1s period before the start of the rTMS burst, using the decibel conversion. The frequency range was normalized by extracting the IAF from the original frequency, and was averaged over IAF ± 1Hz and over the ten left parietal channels (i.e., P7, P5, P3, P1, Pz, PO7, PO3, POz, O1, Oz).

Statistical analysis of the normalized power including ten channels and the entire trial duration from zero to ten seconds was performed for each stimulation intensity separately. First, we used the independent samples t-test to compare rhythmic real and rhythmic sham rTMS protocols in the High-intensity condition. When comparing the real and sham rhythmic protocols, we focused primarily on the high intensity condition because our participants received only one sham rTMS session corresponding to the high intensity condition in the main experiment. Note that in the sham protocol we fixed the stimulation intensity at 29% of the device output. To compare the rhythmic and arrhythmic conditions we used dependent sample t-tests separately for each intensity condition at IAF ± 1 Hz. A non-parametric Monte Carlo approach with 1,000 randomizations was performed to estimate the probability of whether a given amount of significant electrodes (p<0.05) could be expected by chance.

## 3. Results

### 3.1. Short-term aftereffect

First, we focused on analyzing the alpha power change following the rTMS bursts and compared it to the baseline value. In the rhythmic conditions, the analysis revealed no statistically significant differences from baseline in any of the intensity conditions (see Fig 2). Note that in the Medium intensity condition the change was nearly significant (p = 0.07). However, in the arrhythmic conditions there was a significant change with the Medium intensity (p = 0.03), but not with any other intensity (see Fig 2B). Lastly, the analysis revealed that the alpha power did not change significantly from baseline after the sham protocol (Fig 2C). Note that the present study used only one sham condition as a control for the High intensity rhythmic condition.

**Fig 2.**
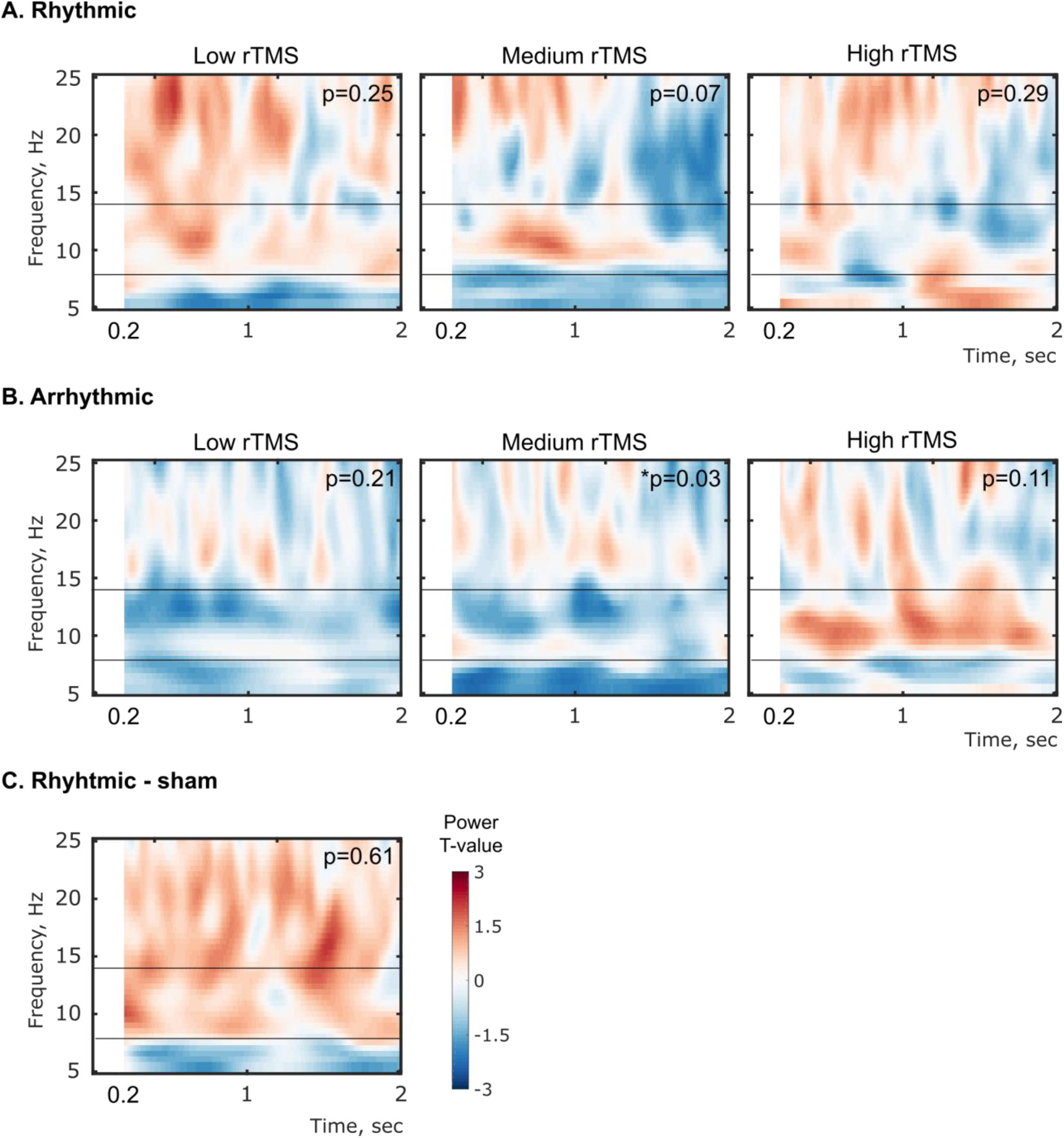
Alpha power change after the rTMS bursts compared with the baseline time period (activation vs. baseline analysis). Time-frequency plots show the power in the range from 5 to 25 Hz (A) in the rhythmic, main, (B) in the arrhythmic, control and (C) in the sham rTMS protocols. Horizontal lines represent the limits of alpha rhythm (8-14 Hz). Zero on the abscissa corresponds to the time of stimulation offset. Statistical analysis was performed with a gap of 200 ms to reduce the influence of residual TMS artifacts.

### 3.2. Long-term aftereffect

In the following analyses, we focused on the IAF, because the entrainment hypothesis predicts that the most pronounced effects should occur in frequencies at and close to the IAF [15]. We compared the rhythmic and sham protocols in the High intensity condition using a non-parametric cluster-based permutation test of the normalized alpha power. The analysis did not reveal any significant difference between the real and sham groups (p = 0.30; Fig 3).

**Fig 3.**
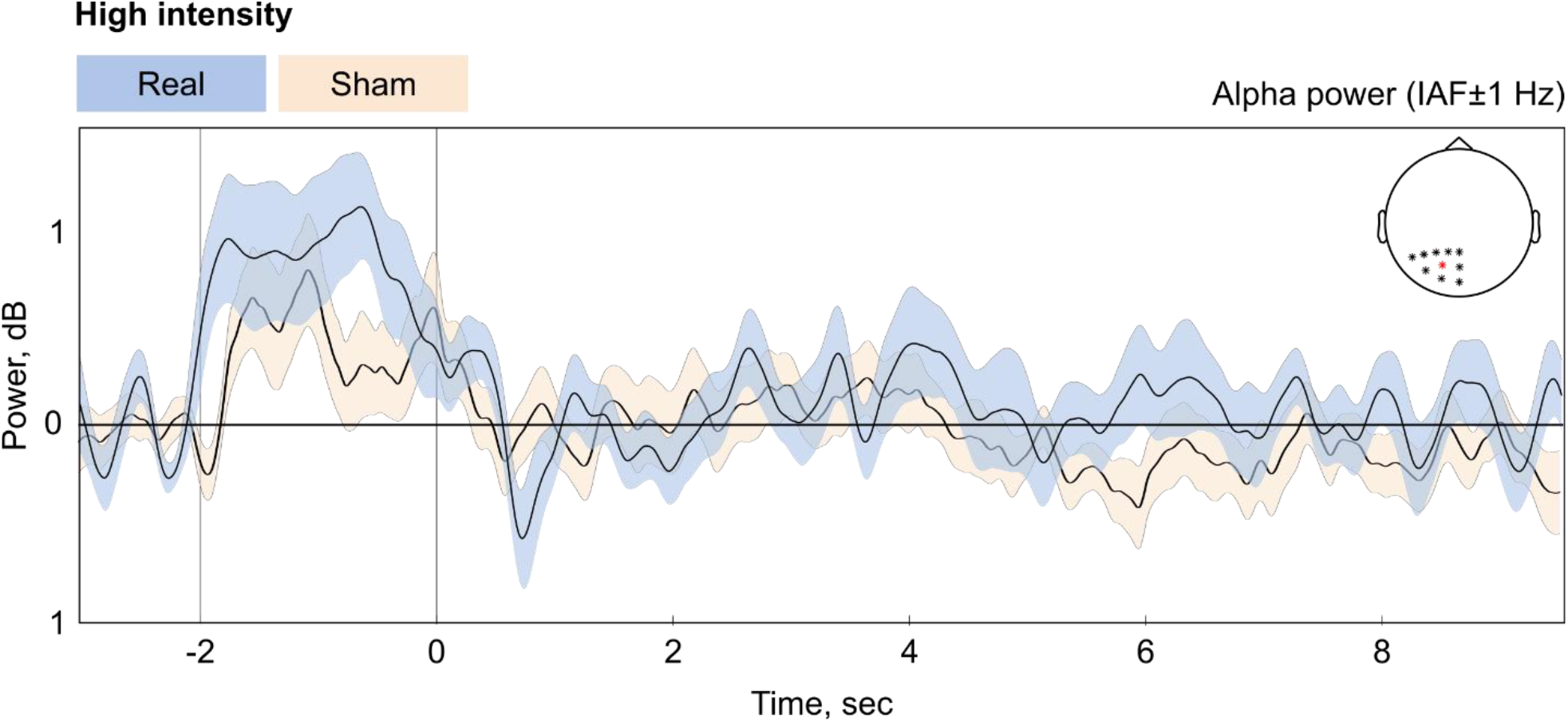
Real rTMS did not change the spectral power relative to the sham rTMS at the individual alpha frequency. The plots show the mean (black line) and SEM (shaded area) of normalized alpha power during the whole trial. The power at IAF ± 1Hz was averaged over ten parietal channels around the stimulation electrode – PO3 (red). The vertical lines at −2 and zero seconds represent stimulation onset and offset, respectively. Note that we aligned the analysis relative to the end of rTMS bursts. Thus, the exact beginning at −2 second varies according to the IAF.

Next, we compared the rhythmic and arrhythmic protocols using non-parametric cluster-based permutation tests on the normalized alpha power. Again, the test revealed no significant differences between these protocols either in the High (p = 0.18), Medium (p = 0.08), or Low (p = 0.23) intensity conditions (see Fig 4).

**Fig 4.**
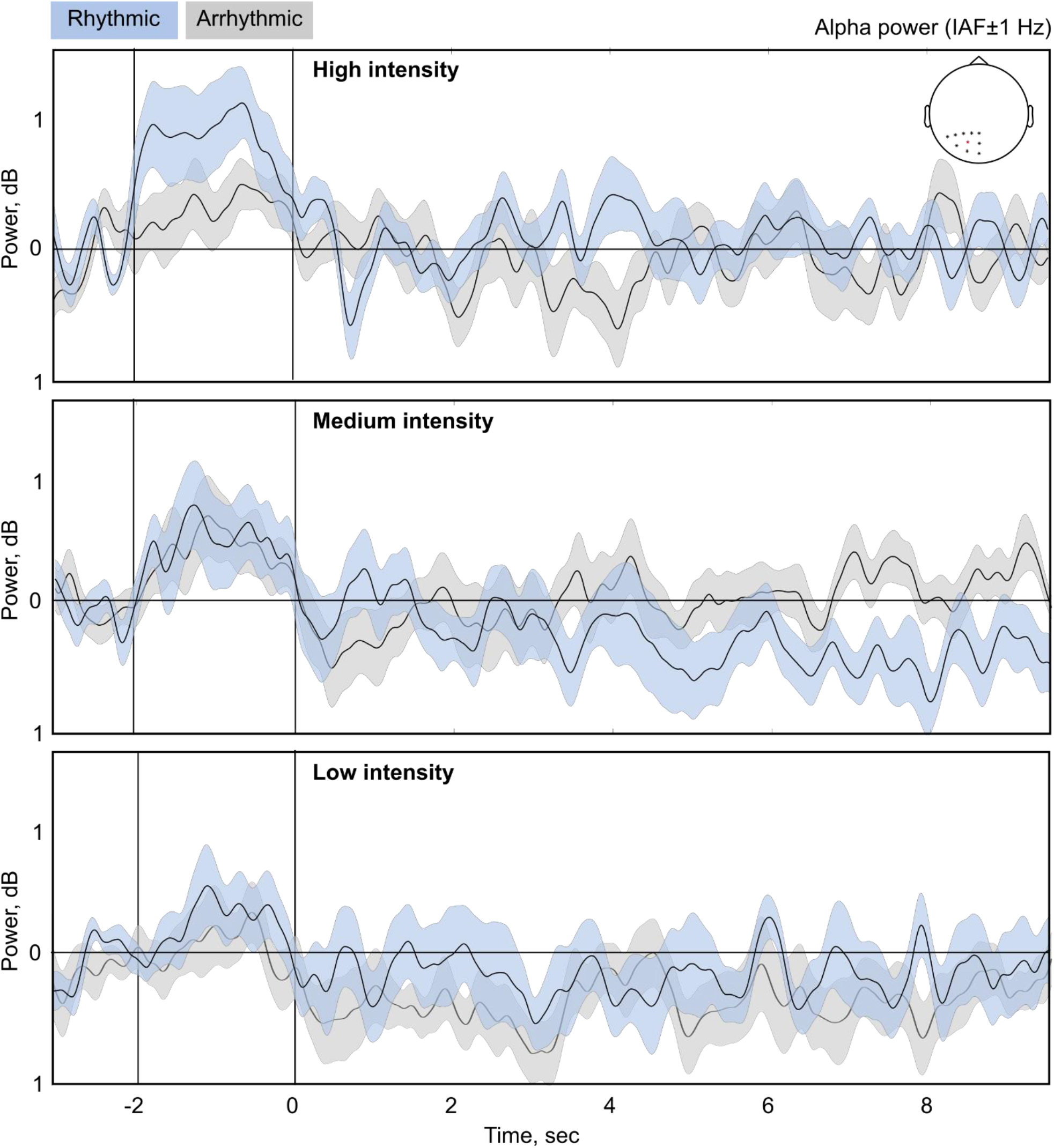
Lack of significant differences in the individual alpha power between rhythmic and arrhythmic rTMS. The plots show the mean (black line) and SEM (shaded area) of alpha power after rTMS bursts (time = 0). The power is normalized to the 1-second-long baseline period directly before the rTMS bursts with decibel correction and averaged over groups and ten parietal channels. Alpha power is extracted at IAF ± 1Hz. Statistical analysis showed no significant difference between the rhythmic and arrhythmic conditions for any stimulation intensity. The vertical lines at −2 and zero seconds represent stimulation onset and offset, respectively. Note that we aligned the analysis relative to the end of rTMS bursts. Thus, the exact beginning at −2 second varies according to the IAF.

These findings indicate that relative to the arrhythmic, control conditions, real rTMS at ca. 20 and 50 mV/mm peak absolute electric field did not change the spectral power in the inter-burst intervals in the individual alpha frequency ± 1 Hz range. There was a non-significant (p = 0.08) decrease in alpha power relative to the arrhythmic condition, real rTMS at ca. 35 mV/mm for up to 10 seconds.

## 4. Discussion

In the present study, we investigated the electrophysiological aftereffects of rhythmic, arrhythmic, and sham rTMS protocols in humans. We defined aftereffects as changes in the alpha power (8-14 Hz) during the inter-burst intervals. We measured short-term aftereffects, i.e. up to two seconds after stimulation, and long-term aftereffects, i.e. from two to ten seconds after stimulation. We expected that rhythmic rTMS would entrain alpha oscillations and lead to increased alpha power after rTMS [7]. Based on the entrainment echo hypothesis, we expected alpha power to be increased for up to ca. two seconds after each burst with rhythmic stimulation. We also expected that neither sham nor arrhythmic rTMS would have any aftereffects on power modulation.

Contrary to our expectations, we observed no aftereffects on alpha power in the rhythmic rTMS protocols with all intensities. In the medium intensity condition, we observed a significant decrease in alpha power in the arrhythmic, and a slight, but non-significant increase in the rhythmic protocol. When studying the entire ten-second inter-burst interval, we found no significant differences in alpha power between the rhythmic and sham or rhythmic and arrhythmic protocols.

### 4.1. Do stronger electric fields induce more robust aftereffects on alpha power?

Compared to conventional rTMS studies that typically use electric fields of ca. 100 mV/mm, the present study applied field strengths that were several times weaker ranging from 20 to 50 mV/mm. One might argue that the applied electric field strength was simply too weak to induce any aftereffects. Following the above argument, one should find more robust aftereffects on alpha power in studies using much stronger stimulation intensities and thus greater electric field strengths. To gain a comprehensive overview, we performed a systematic literature search on rTMS studies using conventional intensities published between 1989 and 2017 (see S1 Appendix for details).

In this search, we focused on studies that evaluated the aftereffects of 10 Hz rTMS on alpha power. We identified 16 eligible articles; ten of which described no aftereffects after rTMS. Two articles described an increase, two articles observed both an increase and a decrease, and one article described a decrease. One article reported incomplete statistical tests to support the claimed aftereffect (e.g., post-hoc tests were missing; see S1. Table for more details). One plausible reason for the contradictory findings may be the known variability in the stimulation parameters, such as the number of pulses, duration of the inter-train intervals, the neuronal state of the stimulated area, etc. [16].

Moreover, these studies also differ in how they operationalize the rTMS-induced aftereffects. Whereas some studies focused on the short inter-burst intervals [e.g., 17], others analyzed the time interval after the end of the rTMS protocol [e.g., 18]. Furthermore, studies may also differ in whether they evaluate the aftereffects directly after the end of the rTMS protocol or after a certain delay period [e.g., 19]. In the present literature search, this delay period varied from several minutes [e.g., 20] up to one week [e.g., 21]. Finally, these studies recruited healthy persons as well as patients (e.g., medication resistant major depression [20]), which is an important factor to consider when evaluating the aftereffects of rTMS.

Taken together, it is difficult to draw comprehensive conclusions about the expected direction of the EEG aftereffects following 10 Hz rTMS. Therefore, the result of the literature analysis was that the evidence about the aftereffects on spectral power in conventional rTMS studies is currently inconclusive.

### 4.2. Outlook and conclusions

At conventional intensities, 10 Hz rTMS is supposed to increase the corticospinal excitability level [16]. The most typical outcome measure in humans is the peak-to-peak amplitude of the single pulse TMS-induced motor evoked potential. Many studies have found increased motor evoked potential amplitudes after the end of a 10 Hz rTMS protocol that lasted for a few minutes [22]. Inhibitory synaptic effects likely play a significant role in the pattern of aftereffects. For instance, a previous *in vitro* tissue culture study provided evidence that 10 Hz repetitive magnetic stimulation induced long-term potentiation in inhibitory synapses [23]. Moreover, scalp EEG alpha oscillations have been associated with cortical inhibition in humans [24]. Therefore, future studies should also investigate the aftereffects of 10 Hz rTMS on the corticospinal excitability level together with the EEG changes when applying weak electric fields, such as in the present study.

In the present study we focused on electrophysiological aftereffect recorded during the inter-burst intervals. At medium intensities (ca. 35 mV/mm), arrhythmic rTMS significantly reduced the alpha power shortly after the rTMS bursts, while the increase in alpha power after rhythmic rTMS was not statistically significant. These findings may be explained by previous observations that cortical inhibitory mechanisms might have lower intensity thresholds than those producing excitation [25]. It remains to be seen which electric field intensities can induce more robust and long-term aftereffects that are manifest for up to several minutes or even longer after the end of the protocol.

## 5. Conflict of interest

We wish to confirm that there are no known conflicts of interest associated with this publication and there has been no significant financial support for this work that could have influenced its outcome.

## 6. Authors contribution

Authors contribution was prepared according to the Contributor Roles Taxonomy. Conceptualization: ZT; Study design: EZ, MM, ZT and WP; Formal analysis: EZ; Funding acquisition: ZT, WP; Investigation: EZ and ZT; Methodology: EZ and ZT; Project administration: EZ, ZT and WP; Software: EZ and ZT; Supervision: MM and WP; Visualization: EZ and ZT; Writing - original draft: EZ and ZT with the critical contribution of all authors.

## 7. Acknowledgement

We thank Jana Thiel for her help in maintaining the EEG system and recruiting the volunteers. We would like to thank Prof. Thomas Crozier for his comments on the manuscript. The study was supported in part by the starting grant of the University Medical Center Göttingen awarded to ZT.

## 9. Supplemental information

### S1 Appendix

We found 194 articles between January 2009 and December 2017 that described studies using rTMS at the alpha frequency band in humans. We selected studies delivering rTMS at 10 Hz and at individualized frequencies at alpha or mu rhythms. We excluded 145 articles that did not use the EEG to evaluate the effects of rTMS. We removed six articles that sequentially combined 1 Hz rTMS with 10 Hz rTMS as well as two prospective clinical trials. We identified 41 articles that combined rTMS with EEG measurements, 17 of which evaluated the effects of rTMS by assessing spectral power. We further excluded four articles that focused on immediate electrophysiological effects. Ten of the remaining thirteen articles used a fixed 10 Hz stimulation frequency. Two articles set the stimulation frequency at the individual mu rhythm, and one at the individual alpha rhythm (see Part I in S1. Table).

We further divided the 13 articles based on the time period in which they analyzed the rTMS-induced electrophysiological aftereffects. High-frequency rTMS (≥ 5 Hz) protocols deliver the stimulation in short bursts/trains and therefore employ several seconds of inter-train intervals between each burst. For example, one can deliver 1,000 rTMS pulses in 20 bursts, using 50 pulses in each burst and 25 s inter-train intervals. The role of the inter-train interval is at least twofold: they prevent coil overheating, and are important for patient safety. Without inter-train intervals, the likelihood increases that high-frequency rTMS might induce an epileptic seizure even in healthy individuals. The short inter-train interval also allows recording and analyzing simultaneous scalp EEG periods that are free of rTMS-induced artifacts. Therefore, the EEG analysis can focus on these short inter-train intervals. It can start directly after the last pulse or several minutes after the end of the protocol. We identified four articles that analyzed the aftereffects during the inter-train intervals. Six articles focused on aftereffects occurring directly after the last pulse and four after the end of the stimulation protocol. This latter period varied between several minutes to one week.

**S1. Table.**
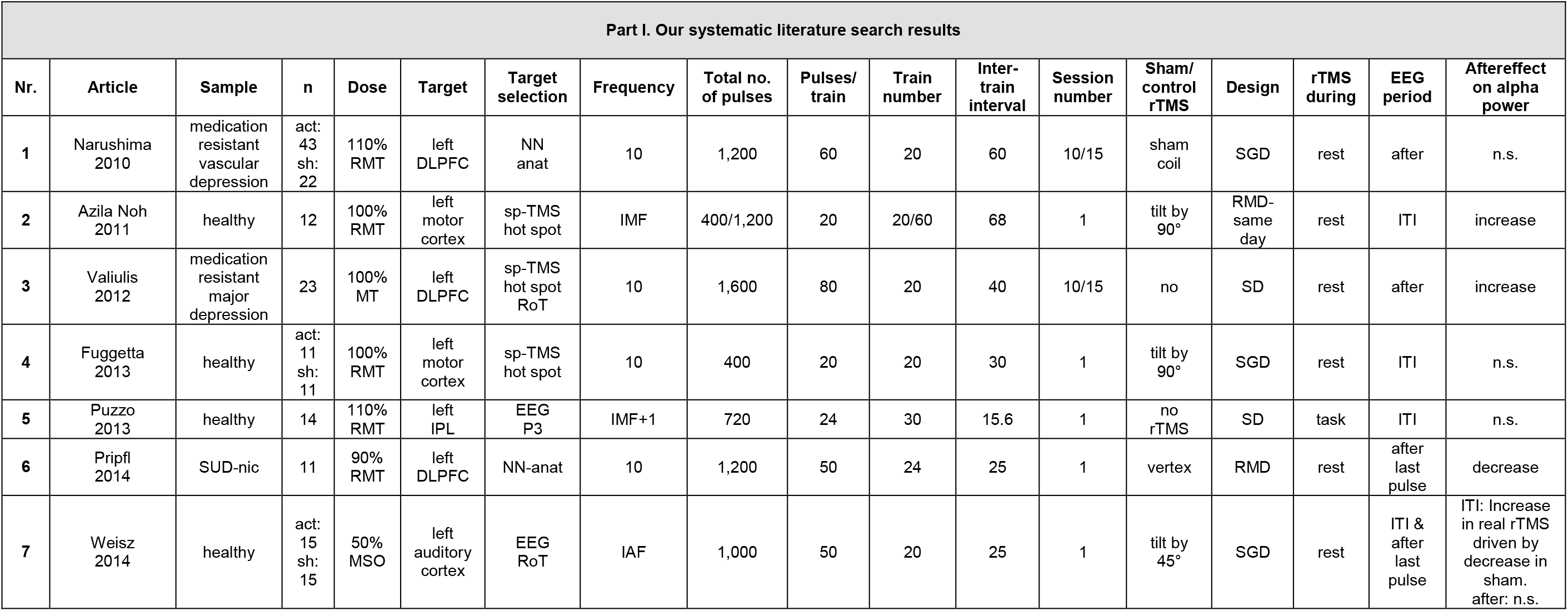
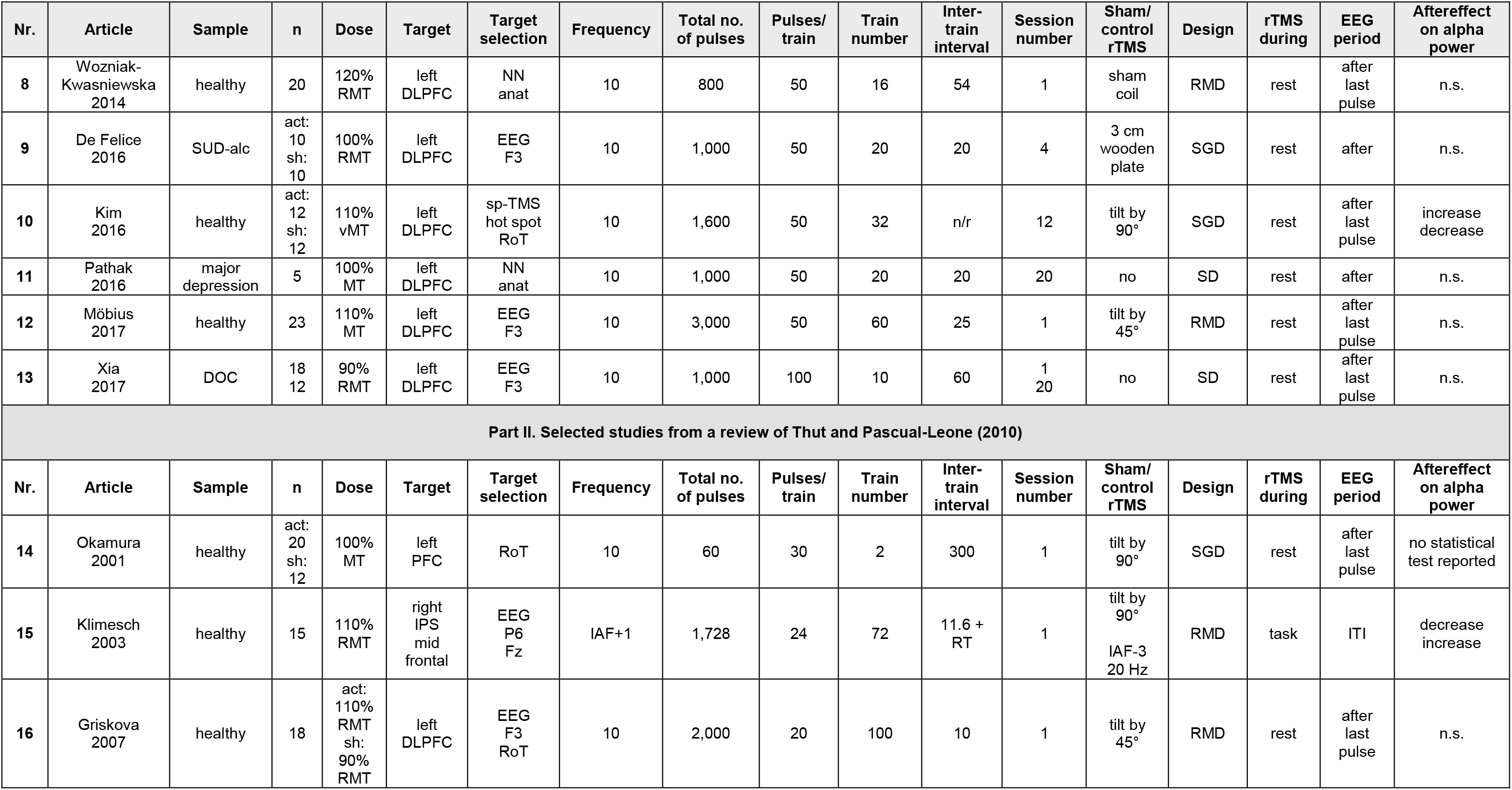
Summary of studies investigating the rTMS-induced electrophysiological aftereffects. Abbreviations: act: active/real stimulation; DLPFC: dorso-lateral prefrontal cortex; EEG RoT: EEG electrode landmark and rule of thumb; IAF: individual alpha frequency; IMF: individual mu frequency; IPL: inferior parietal lobe; iIPS: inferior intraparietal sulcus; ITI: inter-train intervals; MT: motor threshold; NN anat: neuronavigation based on individual anatomy; n: sample size; No.: number; n.s.: not significant; PT: phosphene threshold; RMD: repeated measures design; RMT: resting motor threshold; RoT: rule of thumb; RT: participants’ reaction time; S1: primary somatosensory cortex; SD: single design without sham or control rTMS; sh: sham stimulation; SGD: separate group design; SPL: superior parietal lobule; sp-TMS: single pulse TMS; vMT: visual motor threshold.

| Part I. Our systematic literature search results |  |  |  |  |  |  |  |  |  |  |  |  |  |  |  |  |  |
| --- | --- | --- | --- | --- | --- | --- | --- | --- | --- | --- | --- | --- | --- | --- | --- | --- | --- |
| Nr. | Article | Sample | n | Dose | Target | Target selection | Frequency | Total no. of pulses | Pulses/train | Train number | Inter-train interval | Session number | Sham/control rTMS | Design | rTMS during | EEG period | Aftereffect on alpha power |
| 1 | Narushima 2010 | medication resistant vascular depression | act: 43<br>sh: 22 | 110% RMT | left DLPFC | NN anat | 10 | 1,200 | 60 | 20 | 60 | 10/15 | sham coil | SGD | rest | after | n.s. |
| 2 | Azila Noh 2011 | healthy | 12 | 100% RMT | left motor cortex | sp-TMS hot spot | IMF | 400/1,200 | 20 | 20/60 | 68 | 1 | tilt by 90° | RMD-same day | rest | ITI | increase |
| 3 | Valiulis 2012 | medication resistant major depression | 23 | 100% MT | left DLPFC | sp-TMS hot spot RoT | 10 | 1,600 | 80 | 20 | 40 | 10/15 | no | SD | rest | after | increase |
| 4 | Fuggetta 2013 | healthy | act: 11<br>sh: 11 | 100% RMT | left motor cortex | sp-TMS hot spot | 10 | 400 | 20 | 20 | 30 | 1 | tilt by 90° | SGD | rest | ITI | n.s. |
| 5 | Puzzo 2013 | healthy | 14 | 110% RMT | left IPL | EEG P3 | IMF+1 | 720 | 24 | 30 | 15.6 | 1 | no rTMS | SD | task | ITI | n.s. |
| 6 | Pripfl 2014 | SUD-nic | 11 | 90% RMT | left DLPFC | NN-anat | 10 | 1,200 | 50 | 24 | 25 | 1 | vertex | RMD | rest | after last pulse | decrease |
| 7 | Weisz 2014 | healthy | act: 15<br>sh: 15 | 50% MSO | left auditory cortex | EEG RoT | IAF | 1,000 | 50 | 20 | 25 | 1 | tilt by 45° | SGD | rest | ITI & after last pulse | ITI: Increase in real rTMS driven by decrease in sham. after: n.s. |

| Nr. | Article | Sample | n | Dose | Target | Target selection | Frequency | Total no. of pulses | Pulses/ train | Train number | Inter-train interval | Session number | Sham/ control rTMS | Design | rTMS during | EEG period | Aftereffect on alpha power |
| --- | --- | --- | --- | --- | --- | --- | --- | --- | --- | --- | --- | --- | --- | --- | --- | --- | --- |
| 8 | Wozniak-Kwasniewska 2014 | healthy | 20 | 120% RMT | left DLPFC | NN anat | 10 | 800 | 50 | 16 | 54 | 1 | sham coil | RMD | rest | after last pulse | n.s. |
| 9 | De Felice 2016 | SUD-alc | act: 10<br>sh: 10 | 100% RMT | left DLPFC | EEG F3 | 10 | 1,000 | 50 | 20 | 20 | 4 | 3 cm wooden plate | SGD | rest | after | n.s. |
| 10 | Kim 2016 | healthy | act: 12<br>sh: 12 | 110% vMT | left DLPFC | sp-TMS hot spot RoT | 10 | 1,600 | 50 | 32 | n/r | 12 | tilt by 90° | SGD | rest | after last pulse | increase decrease |
| 11 | Pathak 2016 | major depression | 5 | 100% MT | left DLPFC | NN anat | 10 | 1,000 | 50 | 20 | 20 | 20 | no | SD | rest | after | n.s. |
| 12 | Möbius 2017 | healthy | 23 | 110% MT | left DLPFC | EEG F3 | 10 | 3,000 | 50 | 60 | 25 | 1 | tilt by 45° | RMD | rest | after last pulse | n.s. |
| 13 | Xia 2017 | DOC | 18<br>12 | 90% RMT | left DLPFC | EEG F3 | 10 | 1,000 | 100 | 10 | 60 | 1<br>20 | no | SD | rest | after last pulse | n.s. |
| Part II. Selected studies from a review of Thut and Pascual-Leone (2010) |  |  |  |  |  |  |  |  |  |  |  |  |  |  |  |  |  |
| Nr. | Article | Sample | n | Dose | Target | Target selection | Frequency | Total no. of pulses | Pulses/ train | Train number | Inter-train interval | Session number | Sham/ control rTMS | Design | rTMS during | EEG period | Aftereffect on alpha power |
| 14 | Okamura 2001 | healthy | act: 20<br>sh: 12 | 100% MT | left PFC | RoT | 10 | 60 | 30 | 2 | 300 | 1 | tilt by 90° | SGD | rest | after last pulse | no statistical test reported |
| 15 | Klimesch 2003 | healthy | 15 | 110% RMT | right IPS mid frontal | EEG P6 Fz | IAF+1 | 1,728 | 24 | 72 | 11.6 + RT | 1 | tilt by 90°<br>IAF-3 20 Hz | RMD | task | ITI | decrease increase |
| 16 | Griskova 2007 | healthy | 18 | act: 110% RMT<br>sh: 90% RMT | left DLPFC | EEG F3 RoT | 10 | 2,000 | 20 | 100 | 10 | 1 | tilt by 45° | RMD | rest | after last pulse | n.s. |

## Notes

### Competing Interest Statement

The authors have declared no competing interest.

## References

[1] Anastassiou CA, Perin R, Markram H, Koch C. Ephaptic coupling of cortical neurons. Nat Neurosci 2011;14:217–23. https://doi.org/10.1038/nn.2727.

[2] Paulus W, Peterchev AV, Ridding M. Transcranial electric and magnetic stimulation: technique and paradigms. Handb Clin Neurol 2013;116:329–42.

[3] Lenz M, Vlachos A. Releasing the Cortical Brake by Non-Invasive Electromagnetic Stimulation ? rTMS Induces LTD of GABAergic Neurotransmission. Front Neural Circuits 2016;10:96. https://doi.org/10.3389/fncir.2016.00096.

[4] Zmeykina E, Mittner M, Paulus W, Turi Z. Weak rTMS-induced electric fields produce neural entrainment in humans. Sci Rep 2020;10:1–16. https://doi.org/10.1038/s41598-020-68687-8.

[5] Ilmoniemi RJ, Hernandez-Pavon JC, Makela NN, Metsomaa J, Mutanen TP, Stenroos M, et al. Dealing with artifacts in TMS-evoked EEG. Proc Annu Int Conf IEEE Eng Med Biol Soc 2015:230–3. https://doi.org/10.1109/EMBC.2015.7318342.

[6] Thut G, Pascual-Leone A. A Review of Combined TMS-EEG Studies to Characterize Lasting Effects of Repetitive TMS and Assess Their Usefulness in Cognitive and Clinical Neuroscience. Brain Topogr 2010;22:219–32. https://doi.org/10.1007/s10548-009-0115-4.

[7] Hanslmayr S, Matuschek J, Fellner MC. Entrainment of prefrontal beta oscillations induces an endogenous echo and impairs memory formation. Curr Biol 2014;24:904–9. https://doi.org/10.1016/j.cub.2014.03.007.

[8] Oldfield RC. The assessment and analysis of handedness: The Edinburgh inventory. Neuropsychologia 1971;9:97–113. https://doi.org/10.1016/0028-3932(71)90067-4.

[9] Thielscher A, Antunes A, Saturnino GB. Field modeling for transcranial magnetic stimulation: A useful tool to understand the physiological effects of TMS? Proc Annu Int Conf IEEE Eng Med Biol Soc EMBS 2015:222–5. https://doi.org/10.1109/EMBC.2015.7318340.

[10] Opitz A, Windhoff M, Heidemann RM, Turner R, Thielscher A. How the brain tissue shapes the electric field induced by transcranial magnetic stimulation. Neuroimage 2011;58:849–59. https://doi.org/10.1016/j.neuroimage.2011.06.069.

[11] Romei V, Thut G, Mok RM, Schyns PG, Driver J. Causal implication by rhythmic transcranial magnetic stimulation of alpha frequency in feature-based local vs. global attention. Eur J Neurosci 2012;35:968–74. https://doi.org/10.1111/j.1460-9568.2012.08020.x.

[12] Thut G, Schyns PG, Gross J. Entrainment of perceptually relevant brain oscillations by non ‑ invasive rhythmic stimulation of the human brain. Front Psychol 2011;2:170. https://doi.org/10.3389/fpsyg.2011.00170.

[13] Albouy P, Weiss A, Baillet S, Zatorre RJ. Selective Entrainment of Theta Oscillations in the Dorsal Stream Causally Enhances Auditory Working Memory Performance. Neuron 2017;94:193–206.e5. https://doi.org/10.1016/j.neuron.2017.03.015.

[14] Wu W, Keller CJ, Rogasch NC, Longwell P, Shpigel E, Rolle CE, et al. ARTIST: A fully automated artifact rejection algorithm for single-pulse TMS-EEG data. Hum Brain Mapp 2018;39:1607–25. https://doi.org/10.1002/hbm.23938.

[15] Glass L. Synchronization and rhythmic processes in physiology. Naure 2001;410:277–84.

[16] Huang Y-Z, Lu M-K, Antal A, Classen J, Nitsche M, Ziemann U, et al. Plasticity induced by non-invasive transcranial brain stimulation: A position paper. Clin Neurophysiol 2017;128:2318–29. https://doi.org/10.1016/j.clinph.2017.09.007.

[17] Puzzo I, Cooper NR, Cantarella S, Fitzgerald PB, Russo R. The effect of rTMS over the inferior parietal lobule on EEG sensorimotor reactivity differs according to self-reported traits of autism in typically developing individuals $. Brain Res 2013;1541:33–41. https://doi.org/10.1016/j.brainres.2013.10.016.

[18] Woźniak-Kwaśniewska A, Szekely D, Aussedat P, Bougerol T, David O. Changes of oscillatory brain activity induced by repetitive transcranial magnetic stimulation of the left dorsolateral prefrontal cortex in healthy subjects. Neuroimage 2014;88:91–9. https://doi.org/10.1016/j.neuroimage.2013.11.029.

[19] Weisz N, Lüchinger C, Thut G, Müller N. Effects of individual alpha rTMS applied to the auditory cortex and its implications for the treatment of chronic tinnitus. Hum Brain Mapp 2014;35:14–29. https://doi.org/10.1002/hbm.22152.

[20] Valiulis V, Gerulskis G, Dapšys K, Vištartaitė G, Šiurkutė A. Electrophysiological differences between high and low frequency rTMS protocols in depression treatment. Acta Neurobiol Exp (Wars) 2012;72:283–95.

[21] Narushima K, Mccormick LM, Yamada T, Thatcher RW, Robinson RG. Subgenual Cingulate Theta Activity Predicts Treatment Response of Repetitive Transcranial Magnetic Stimulation in Participants With Vascular Depression. J Neuropsychiatry Clin Neurosci 2010;22:75–84.

[22] Arai N, Okabe S, Furubayashi T, Mochizuki H, Iwata NK, Hanajima R, et al. Differences in after-effect between monophasic and biphasic high-frequency rTMS of the human motor cortex. Clin Neurophysiol 2007;118:2227–33. https://doi.org/10.1016/j.clinph.2007.07.006.

[23] Lenz M, Galanis C, Mu F, Opitz A, Wierenga CJ. Repetitive magnetic stimulation induces plasticity of inhibitory synapses. Nat Commun 2016:7:10020. https://doi.org/10.1038/ncomms10020.

[24] Klimesch W, Sauseng P, Hanslmayr S. EEG alpha oscillations: The inhibition-timing hypothesis. Brain Res Rev 2007;53:63–88. https://doi.org/10.1016/j.brainresrev.2006.06.003.

[25] Moliadze V, Atalay D, Antal A, Paulus W. Close to threshold transcranial electrical stimulation preferentially activates inhibitory networks before switching to excitation with higher intensities. Brain Stimul 2012;5:505–11. https://doi.org/10.1016/j.brs.2011.11.004.

